# ATM and MSH2 control blunt DNA end joining in immunoglobulin class switch recombination

**DOI:** 10.1101/2022.07.28.501875

**Authors:** Emily Sible, Mary Attaway, Giuseppe Fiorica, Genesis Michel, Jayanta Chaudhuri, Bao Q. Vuong

## Abstract

Class switch recombination (CSR) produces secondary immunoglobulin isotypes and requires AID-dependent DNA deamination of intronic switch (S) regions within the immunoglobulin heavy chain (*Igh*) gene locus. Non-canonical repair of deaminated DNA by mismatch repair (MMR) or base excision repair (BER) creates DNA breaks that permit recombination between distal S regions. ATM-dependent phosphorylation of AID at serine-38 (pS38-AID) promotes its interaction with APE1, a BER protein, suggesting that ATM regulates CSR through BER. However, pS38-AID may also function in MMR during CSR, although the mechanism remains unknown. To examine whether ATM modulates BER- and/or MMR-dependent CSR, *Atm^-/-^* mice were bred to mice deficient for the MMR gene *Msh2*. Surprisingly, the predicted Mendelian frequencies of *Atm^-/-^Msh2^-/-^* adult mice were not obtained. To generate ATM and MSH2-deficient B cells, *Atm* was conditionally deleted on an *Msh2^-/-^* background using a floxed ATM allele [*Atm^f^*] and B cell-specific Cre recombinase expression (*CD23-cre*) to produce a deleted ATM allele (*Atm^D^*). As compared to *Atm^D/D^*and *Msh2^-/-^* mice and B cells, *Atm^D/D^Msh2^-/-^* mice and B cells display a reduced CSR phenotype. Interestingly, Sμ-Sγ1 junctions from *Atm^D/D^Msh2^-/-^*B cells that were induced to switch to IgG1 *in vitro* showed a significant loss of blunt end joins and an increase in insertions as compared to wildtype, *Atm^D/D^*, or *Msh2^-/-^* B cells. This data indicates that the absence of both ATM and MSH2 blocks non-homologous end joining (NHEJ), leading to inefficient CSR. We propose a model whereby ATM and MSH2 function cooperatively to regulate end-joining during CSR through pS38-AID.

**Summary:** Loss of the DNA repair genes *Atm* and *Msh2* produces a novel synthetic lethality in mice. B cell specific deletion of *Atm* on an *Msh2^-/-^* background reduces Ig CSR and inhibits NHEJ.

## Introduction

B cells recognize and eliminate antigens by producing immunoglobulins (Ig), also known as antibodies. Each Ig is composed of two heavy (IgH) and two light (IgL) chain polypeptides, which are held together by disulfide bonds to form the characteristic “Y” shape structure of the Ig [1]. The N-termini of IgH and IgL comprise the variable (V) region of each polypeptide and together they form the antigen binding site of the Ig, whereas the constant region of IgH imparts the effector function of the Ig.

In a process known as Class Switch Recombination (CSR), mature B cells generate double-strand DNA breaks (DSBs) within the immunoglobulin heavy chain gene locus (*Igh*) to diversify the Ig repertoire through a DNA recombination event and produce new Ig isotypes [2–4]. CSR is mediated by the enzymatic activity of activation-induced cytidine deaminase (AID) [5–7], which converts deoxycytidines into deoxyuridines in the repetitive switch (S) region that precedes each *Igh* constant coding exon [5–7]. Non-canonical repair of these U:G mutations in DNA by either mismatch repair (MMR) or base excision repair (BER) creates staggered DNA breaks that promote recombination between distal S regions [8–15] and expression of alternative *Igh* constant coding exons [16].

During BER, uracil DNA Glycosylase (UNG) generates an abasic site by removing the AID-generated uracil base, allowing for subsequent cleavage of the phosphodiester backbone by apurinic/apyrimadinic endonuclease 1 (APE1) to generate a single-strand DNA break (SSB) [13] [17]. In contrast, MMR utilizes a heterodimer of mutS homologs MSH2/MSH6 to recognize DNA base pair mismatches and recruits the mutL homology 1 (MLH1) and PMS1 homolog 2 (PMS2) heterodimer to cleave the phosphodiester backbone 5’ of the mismatch [6].

Exonuclease-1 (EXO1) degrades the DNA to remove the U:G mismatch [18]. Staggered SSBs in close proximity constitute DSBs, which are converted into recombination events. Mice deficient in MSH2 or UNG show reduced levels of switched isotypes as compared to WT mice [8]. Furthermore, simultaneous loss of both MSH2 and UNG blocks CSR completely, demonstrating the complementary function of MMR and BER in CSR [8].

Activation of BER and MMR during CSR is mediated by the phosphorylation of AID at serine 38 (pS38-AID), which promotes the interaction of AID with APE1 [19]. Mice with a homozygous knock-in mutation of the phosphorylation site (*Aicda^S38A/S38A^*), which changes S38 to an alanine, have significantly reduced CSR [20, 21]. *Aicda^S38A/S38A^* mice with homozygous null mutations in *Ung* or *Msh2* show a CSR phenotype comparable to *Ung^-/-^Msh2^-/-^* mice, indicating a critical role of AID phosphorylation in activating BER and MMR during CSR [22]. Furthermore, AID phosphorylation at S38 requires ataxia telangiectasia mutated (ATM), a phosphatidylinositol 3-kinase-related kinase (PIKK) that responds to DSBs and is required for wildtype (WT) CSR [23, 24]. *Atm^-/-^*mice show reduced levels of pS38-AID and impaired AID/APE1 interaction, suggesting that ATM promotes phosphorylation of AID to induce its interaction with APE1 and drive CSR through BER [19]. However, the mechanism by which ATM signals non-canonical, recombinogenic BER during CSR remains unclear.

After BER- and MMR-dependent DSBs are formed in the recombining S regions, nonhomologous end join (NHEJ) factors ligate the broken ends and promote DNA recombination rather than canonical DSB repair [25, 26]. Notably, the absence of NHEJ factors, such as Ku70, Ku80, XRCC4, or DNA ligase IV, reduces CSR efficiency and increases *Igh* chromosomal translocations [27–29]. Without NHEJ, CSR can be completed through a microhomology-mediated end joining pathway, also known as alternative non-homologous end joining (A-EJ), which is more error prone and less efficient than NHEJ [30, 31]. Recombined S-S junctions from WT B cells show ligations resulting primarily from NHEJ [27, 28]. The majority of these NHEJ-repaired S-S junctions show short stretches of homology, or microhomology, between donor and acceptor S regions of less than 4 nucleotides, and the remaining approximately 30% of S-S junctions contain no homology between the recombining S regions and are characterized as a blunt end join [32]. Recombined S-S regions that have been repaired by A-EJ contain longer stretches of homology, which skew towards 5 nucleotides or more [27] [32], yet the conditions under which A-EJ is favored over NHEJ during CSR remain unclear. *Atm^-/-^* B cells exhibit longer microhomologies in recombined S-S junctions, indicating that ATM is required for NHEJ and A-EJ mediates recombinational repair in the absence of ATM [24, 33]. Interestingly, MSH2 has been implicated in regulating A-EJ during CSR in the absence of NHEJ because Sμ-Sγ3 junctions from B cells lacking MSH2 exhibited no microhomology above 5nt [10]. Here, we demonstrate an epistatic role for *Atm* to *Msh2* in CSR. Loss of MSH2 and ATM in B cells leads to a reduction in CSR concurrent with a block in blunt end S-S junctions, suggesting a role for MSH2 in enforcing NHEJ in the absence of ATM. Additionally, we report a novel synthetic lethality between *Atm* and *Msh2*, which implicates ATM or MSH2 as a potential therapeutic target for cancers driven by mutations in either gene.

## Results

### Mutations in *Atm* and *Msh2* induce a developmental lethality late in gestation to early postnatal

BER and MMR function redundantly to direct CSR in B cells [8]. ATM regulates the phosphorylation of AID (p-AID) and its interaction with APE1, a BER protein [19]. To determine the role of ATM in BER-dependent CSR, we bred *Atm^-/-^* and *Msh2^-/-^* mice to generate *Atm^-/-^Msh2^-/-^*mice. *Atm^-/-^* mice carry a PGKneo modified exon at position 5790 in the *Atm* locus that recapitulates mutations observed in ataxia telangiectasia patients [34]. *Msh2^-/-^* mice contain a targeted disruption 5’ of the exon encoding the MSH2 ATP binding domain, resulting in a frameshift and mRNA decay [35]. *Atm^-/-^* and *Msh2^-/-^*single knockout mice survive into adulthood, albeit with mild immune defects and a predisposition to lymphomagenesis [36, 37].

Surprisingly, of the 247 mice genotyped at post-natal day 28 (P28), no *Atm^-/-^Msh2^-/-^* were found **(Table 1**, p=3.3x10^-4^), unveiling a novel synthetic lethal relationship. To determine the developmental stage at which the lethality occurs, embryos were collected from timed mating two weeks after conception (E13.5) and genotyped. Mendelian ratios of E13.5 embryos from a dihybrid cross were obtained (**Table 2**). Newborn pups (P0) from *Atm^+/-^Msh2^-/-^* x *Atm^+/-^Msh2^+/-^*crosses yielded less than half of the expected number of P0 *Atm^-/-^Msh2^-/-^*neonates; however, these findings are not statistically significant (**Table 3**; p = 0.292). Collectively, these data indicate that the *Atm^-/-^Msh2^-/-^*lethality occurs between E13.5 and P0 with an incomplete penetrance of the synthetic lethal phenotype.

**Table 1:** Significant absence of Atm^-/-^Msh2^-/-^ adult mice. The numbers of expected and observed progeny at post-natal day 28 (P28) from a dihybrid cross. Observed progeny were genotyped at approximately four weeks after birth (P28) and represent ≥ 15 litters. P-value determined by χ^2^ test.

| Cross | N | Genotype | Expected | Observed | $\chi^2$<br>p-value |
| --- | --- | --- | --- | --- | --- |
| $Atm^{+/-} Msh2^{+/-}$<br>x<br>$Atm^{+/-} Msh2^{+/-}$ | 247 | $Atm^{+/+} Msh2^{+/+}$<br>$Atm^{+/+} Msh2^{+/-}$<br>$Atm^{+/-} Msh2^{+/+}$<br>$Atm^{+/+} Msh2^{-/-}$<br>$Atm^{+/-} Msh2^{+/-}$<br>$Atm^{+/-} Msh2^{-/-}$<br>$Atm^{-/-} Msh2^{+/+}$<br>$Atm^{-/-} Msh2^{+/-}$<br>$Atm^{-/-} Msh2^{-/-}$ | 15.44<br>30.88<br>30.88<br>15.44<br>61.75<br>30.88<br>15.44<br>30.88<br>15.44 | 22<br>33<br>37<br>18<br>75<br>27<br>17<br>18<br>0 | $3.3 \times 10^{-4}$ |

**Table 2:**
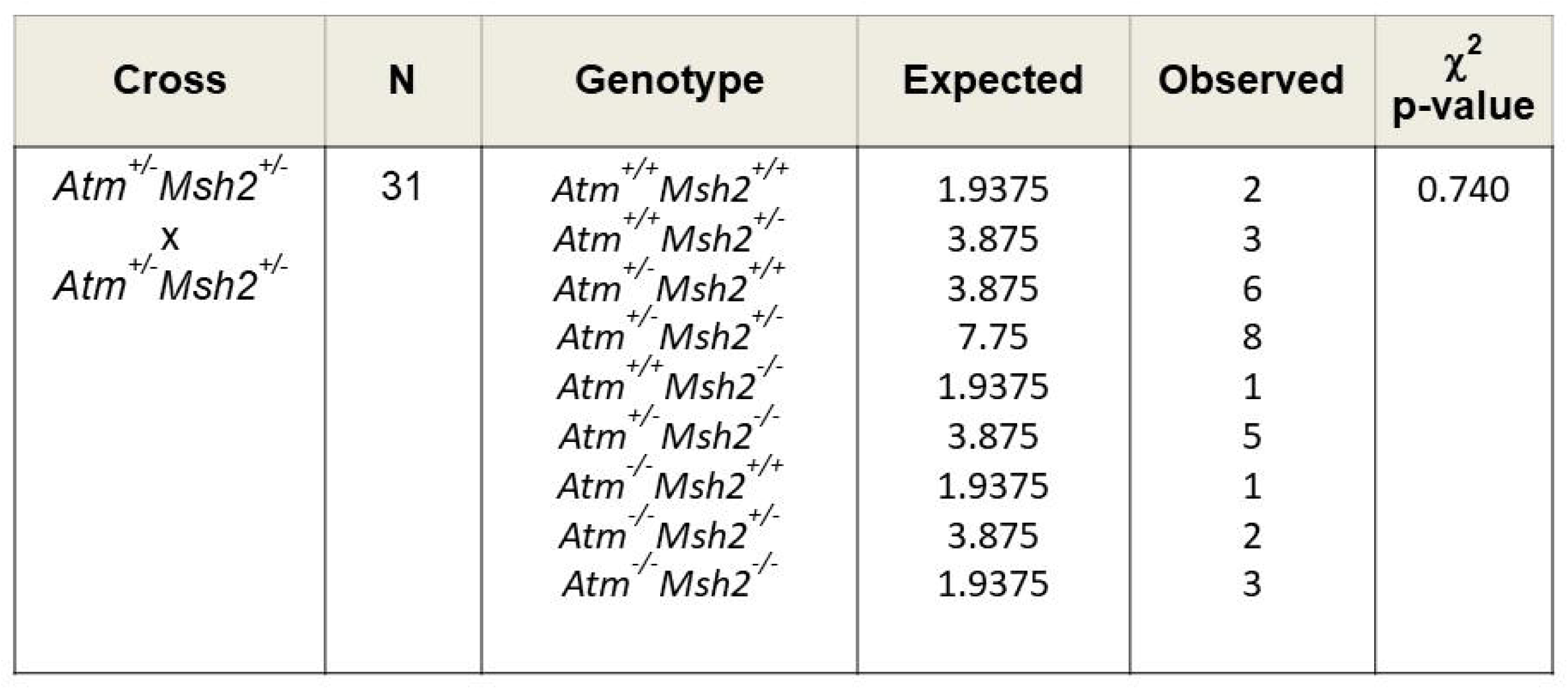
Atm^-/-^Msh2^-/-^ embryos obtained at mid-gestation. The numbers of expected and observed progeny at embryonic day 13.5 (E13.5) that were obtained in a dihybrid cross from timed matings. Observed progeny were obtained from ≥ 3 litters. P-value determined by χ^2^ test.

**Table 3:** Reduced number of *Atm^-/-^Msh2^-/-^* neonates. The numbers of expected and observed newborn progeny (P0) that were obtained from *Atm^+/-^Msh2^+/-^*X *Atm^+/-^Msh2^-/-^*crosses in ≥ 3 litters. P-value determined by χ*^2^* test.

| Cross | N | Genotype | Expected | Observed | $\chi^2$<br>p-value |
| --- | --- | --- | --- | --- | --- |
| $Atm^{+/-} Msh2^{+/-}$<br>x<br>$Atm^{+/-} Msh2^{-/-}$ | 61 | $Atm^{+/+} Msh2^{-/-}$<br>$Atm^{+/+} Msh2^{+/-}$<br>$Atm^{+/-} Msh2^{+/-}$<br>$Atm^{+/-} Msh2^{-/-}$<br>$Atm^{-/-} Msh2^{+/-}$<br>$Atm^{-/-} Msh2^{-/-}$ | 7.625<br>7.625<br>15.25<br>15.25<br>7.625<br>7.625 | 9<br>9<br>19<br>13<br>9<br>2 | 0.292 |

CSR is significantly reduced but not ablated in *Atm^D/D^Msh2^-/-^*mice

To examine the function of ATM and MSH2 in CSR, we bred *Msh2^-/-^*mice to mice that carry a conditional null *Atm* allele (*Atm^f/f^*) and transgenic mice expressing Cre recombinase under the control of a B cell specific promoter (*CD23-Cre^+^*) [38, 39]. The *CD23-Cre^+^Atm^f/f^Msh2^-/-^,* which carry a conditional deletion of *Atm* in B cells and a germline deletion of *Msh2*, are hereafter referred to as *Atm^D/D^Msh2^-/-^.* Unlike *Atm^-/-^Msh2^-/-^* mice, *Atm^D/D^Msh2^-/-^*mice were obtained at the expected Mendelian frequencies (**Supplemental Table 1**), suggesting that the loss of ATM and MSH2 in B cells does not contribute to the *Atm^-/-^Msh2^-/-^*lethality. To investigate the loss of ATM and MSH2 on Ig production, *Atm^D/D^Msh2^-/-^*and control mice were immunized with 4-hydroxy-3-nitrophenylacetyl conjugated to chicken gamma globulin (NP-CGG), a T cell dependent antigen [40, 41]. Serum Igs were analyzed by ELISA prior to and 21 days after a primary and booster immunization of NP-CGG. *Atm^-/-^*, *Aicda^-/-^*, *Msh2^-/-^*, and WT mice showed IgM and IgG1 titers that were consistent with previously published reports (**Figure 1**) [22, 24, 36, 42]. *Atm^D/D^Msh2^-/-^* IgM titers pre- and post-immunization did not differ significantly from WT, *Atm^-/-^, Msh2^-/-^, Atm^D/D^, Msh2^-/-^*, and *CD23-Cre+* mice (**Figure 1**). Given that mature B cells produce IgM without undergoing CSR, these data suggest that *Atm^D/D^Msh2^-/-^*B cells do not have any defects in plasma cell development, Ig production, or Ig secretion.

**Figure 1:**
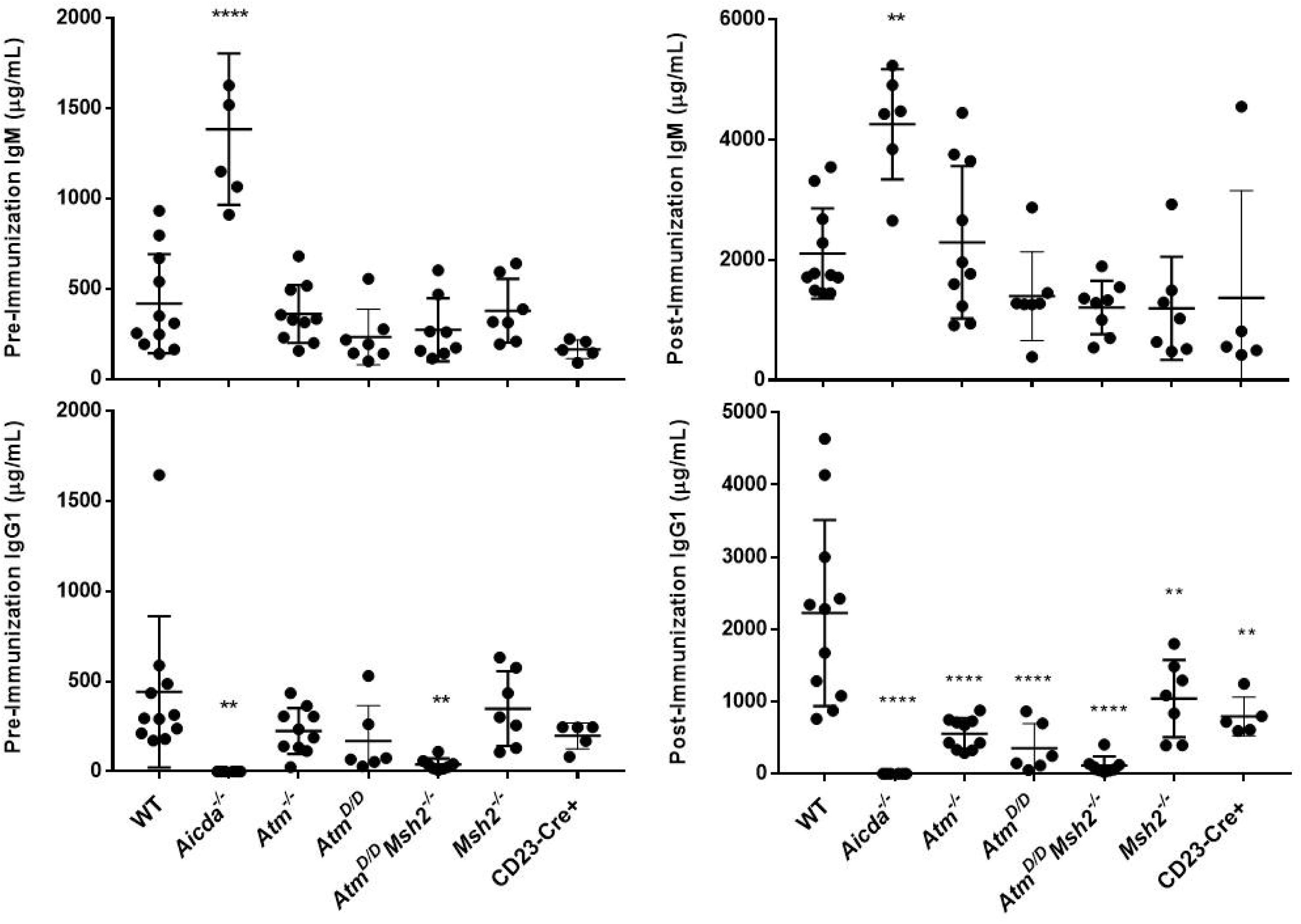
A*t*mD*^/D^Msh2^-/-^*mice show significantly reduced serum IgG1 titers. Serum titers of IgM and IgG1 were assayed before or after immunization with NP-CGG. Statistical analysis was conducted by one way ANOVA for n ≥ 5 (* p<0.05, ** p<0.005, ***p<0.0001, **** p<0.00001). The error bars indicate mean and standard deviation.

To determine whether ATM and MSH2 are required for switch Ig isotypes, IgG1 serum concentrations were quantified before immunization and 21 days after a primary and booster immunization with NP-CGG (**Figure 1**). Consistent with previous reports, IgG1 is absent in *Aicda^-/-^* mice and significantly reduced in *Msh2^-/-^* mice (average 1041 μg/ml) [36, 42]. *Atm^-/-^*mice exhibit a more pronounced reduction in IgG1 (average 554 μg/ml) as compared to *Msh2^-/-^*mice [13, 14]. The average IgG1 titer of *Atm^D/D^*mice (354 μg/ml) did not differ significantly from *Atm^-/-^* mice pre- or post-immunization, suggesting that the conditional deletion of *Atm* effectively recapitulates the IgG1 phenotype observed in the germline deletion of *Atm*. To determine if the reduced IgG1 resulted from defective germinal center B cell (GCBC) development, Peyer’s patches from NP-CGG immunized mice were stained with anti-B220 antibody and PNA and analyzed by flow cytometry (**Supplemental Figure 1**). *Atm^D/D^Msh2^-/-^* mice showed comparable GCBCs numbers as compared to WT, *Atm^D/D^*, and *Msh2^-/-^* mice, suggesting that the reduced IgG1 in *Atm^D/D^Msh2^-/-^* was not a result of defective GC formation but rather a B cell dysfunction in CSR.

To investigate whether *Atm^D/D^Msh2^-/-^* B cells contain an intrinsic defect in CSR, *Atm^D/D^Msh2^-/-^* and control splenic B cells were purified and stimulated *in vitro* with LPS+IL4 to induce CSR to IgG1. Consistent with previous reports [4, 23, 36], CSR to IgG1 is significantly reduced in *Atm^-/-^*and *Msh2*^-/-^ B cells (**Figure 2**). Although trace amounts of ATM protein are detectable by immunoblot in *Atm^D/D^* B cells (**Supplemental Figure 2**), *Atm^D/D^* and *Atm^-/^*^-^ B cells show no significant difference in CSR to IgG1 (p=0.990). Furthermore, *CD23-Cre^+^* B cells have no significant difference in IgG1 CSR as compared to WT cells (p=0.999), demonstrating that the defect in CSR is not due to aberrant B cell specific expression of Cre recombinase. As observed in the serum IgG1 of *Atm^D/D^Msh2^-/-^* mice post-immunization (**Figure 1**), *Atm^D/D^Msh2^-/-^*B cells exhibit significantly reduced, but not ablated, IgG1 (**Figure 2**, p <0.0001). *Atm^D/D^Msh2^-/-^*mouse IgG3 serum titers and *Atm^D/D^Msh2^-/-^* B cells that were stimulated to switch to IgG3 were similarly reduced (**Supplemental Figure 3**), indicating that the defects in CSR in *Atm^D/D^Msh2^-/-^*mice and B cells is not isotype specific. To determine if the reduction in IgG1 was due to reduced proliferation, *Atm^D/D^Msh2^-/-^* or control B cells were labeled with the far red dye SNARF 24 hours after purification and monitored every 24 hours thereafter for dilution of the dye via flow cytometry. No significant differences in cell proliferation were observed between WT, *Atm^D/D^, Msh2^-/-^,* and *Atm^D/D^Msh2^-/-^* B cells (**Supplemental Figure 4**). *CD23-Cre*^+^ B cells diluted the SNARF dye quicker than WT B cells, indicating faster proliferation of *CD23-Cre^+^* B cells (**Supplemental Figure 4**). To examine if the reduction in IgG1 in *Atm^D/D^Msh2^-/-^* B cells was due to defects in germline transcription at the recombining S regions [43], germline transcripts (GLT) for μ and γ1 were quantified from LPS+IL4-stimulated *Atm^D/D^Msh2^-/-^* or control B cells. No significant difference in μ and γ1 germline transcripts was observed across genotypes (**Supplemental Figure 5**), suggesting that the reduction in CSR observed in *Atm^D/D^Msh2^-/-^*B cells is not due to transcriptional defects. Furthermore, immunoblots for AID showed that *Atm^D/D^Msh2^-/-^* B cells expressed AID comparably to WT, *Atm^D/D^*, and *Msh2^-/-^* B cells (**Supplemental Figure 2**), indicating that the CSR defect in *Atm^D/D^Msh2^-/-^* B cells is not due to the absence of AID. In sum, the reduced CSR in *Atm^D/D^Msh2^-/-^* B cells likely results from defects in the molecular pathways regulating the formation or repair of DSBs in S regions rather than in plasma cell development, B cell proliferation, transcriptional activation of recombining S regions, or AID expression.

**Figure 2:**
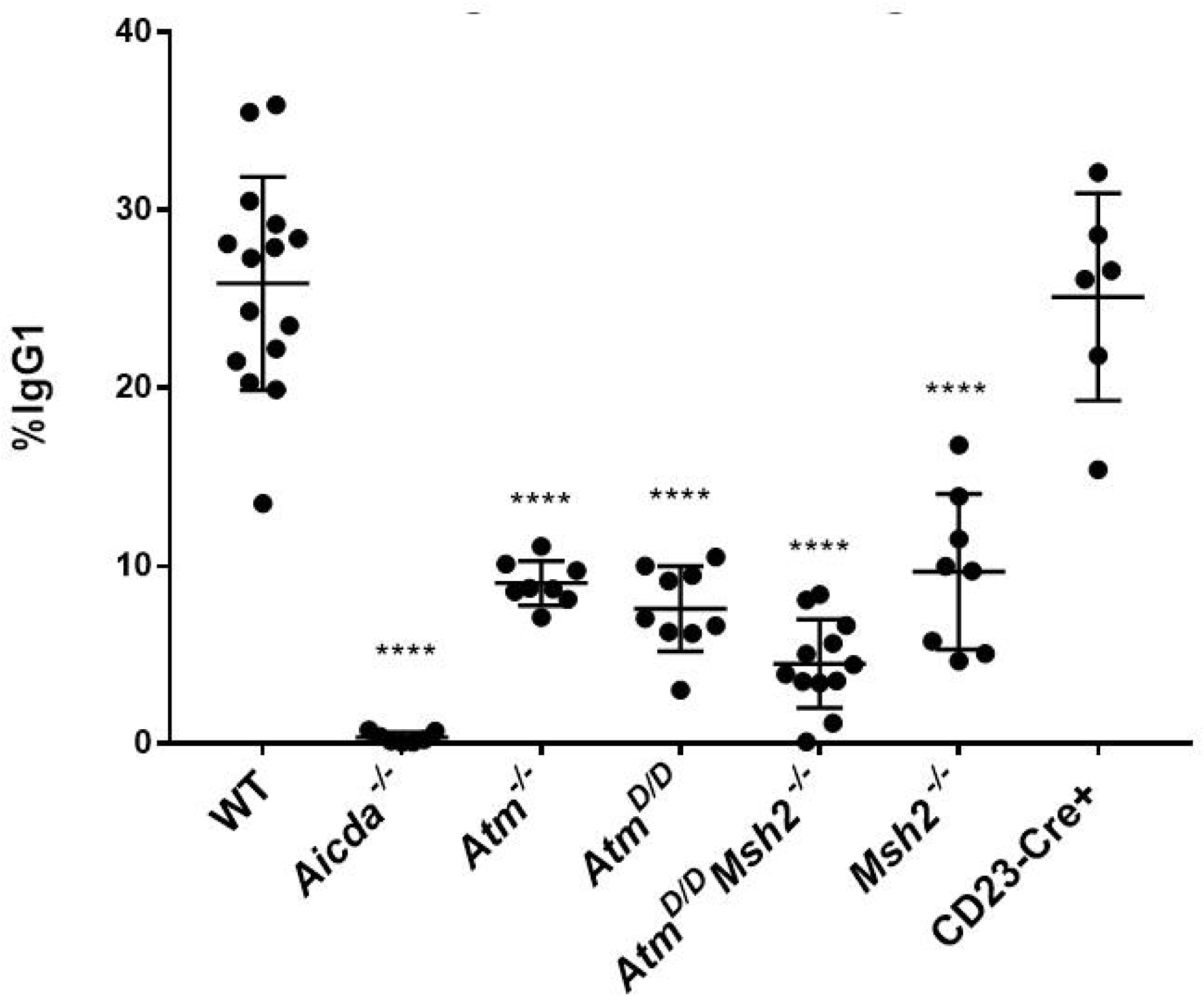
A*t*mD*^/D^Msh2^-/-^*B cells show significantly reduced CSR *in vitro*. Purified B cells of the indicated genotype were stimulated with LPS+IL4 to induce CSR to IgG1, which was measured by flow cytometry. Statistical analysis was conducted by one way ANOVA for n ≥ 6 (****p<0.0001). The error bars indicate mean and standard deviation.

### *Atm^D/D^Msh2^-/-^* B cells exhibit defects in NHEJ

CSR occurs through an intrachromosomal recombination between repetitive, intronic S regions that precede each constant coding exon, resulting in the expression of a new Ig isotype and deletion of the intervening constant exons [10]. Ligation of the DSBs in the recombining S regions requires an end joining pathway to repair the DSB and complete the recombination. In WT B cells, end-joining during CSR requires primarily NHEJ wherein recombined S-S junctions are ligated with no or a few nucleotides of homology (<4 nucleotides) [44]. However, in the absence of NHEJ, B cells completing CSR can use an alternative end-joining (A-EJ) pathway that requires resection of the broken DNA to generate short regions of 5 or more nucleotides (nt) of homology at the recombined S-S junction [44–46]. Therefore, sequencing the recombined S-S junctions analyzes and quantifies the type of DNA repair and end joining pathway that is used in recombining S regions. During end joining, the activity of ATM coupled with the amount of DNA resection at the site of recombination determines whether or not NHEJ will be activated [47, 48]. *Atm^-/^*^-^ B cells show a reduction in blunt end joins and an increase in microhomologies larger than 4nt in S-S junctions, suggesting that ATM drives NHEJ during CSR [23]. In contrast, S-S junctions from *Msh2^-/^*^-^ B cells are joined using blunt or limited microhomology, suggesting that MSH2 promotes A-EJ [16].

To determine whether NHEJ or A-EJ functions in the absence of ATM and MSH2, we analyzed Sμ-Sγ1 junctions from WT, *Atm^D/D^*, *Msh2^-/-^*, and *Atm^D/D^Msh2^-/-^* B cells that were stimulated with LPS+IL4 to induce CSR to IgG1 *in vitro* (**Figure 3**). Sμ-Sγ1 junctions from *Msh2^-/^*^-^ B cells contained primarily blunt end joins or short-stretch microhomology, as previously described for Sμ-Sγ3 junctions in *Msh2^-/-^*B cells [11] [16] [49]. Consistent with Sμ-Sγ1 junctions from *Atm^-/-^* B cells [23], Sμ-Sγ1 junctions from *Atm^D/D^* B cells trend towards longer stretches of microhomology and reduced blunt joins as compared to WT B cells (**Figure 3A**), although these trends have not reached statistical significance. Almost 20% of the Sμ-Sγ1 junctions in *Atm^D/D^Msh2^-/-^* B cells have microhomology lengths of 5 nucleotides or greater (**Figure 3A**). Most strikingly, Sμ-Sγ1 junctions from *Atm^D/D^ Msh2^-/-^* B cells show a significant loss in blunt end joins (**Figure 3A** p=0.007), suggesting a severe block in NHEJ in *Atm^D/D^Msh2^-/-^*B cells. Additionally, coupled with the reduction in blunt end joins, *Atm^D/D^Msh2^-/-^* display a stark increase in insertions at Sμ-Sγ1 junctions as compared to WT B cells (**Figure 3A**, p=0.021).

**Figure 3:**
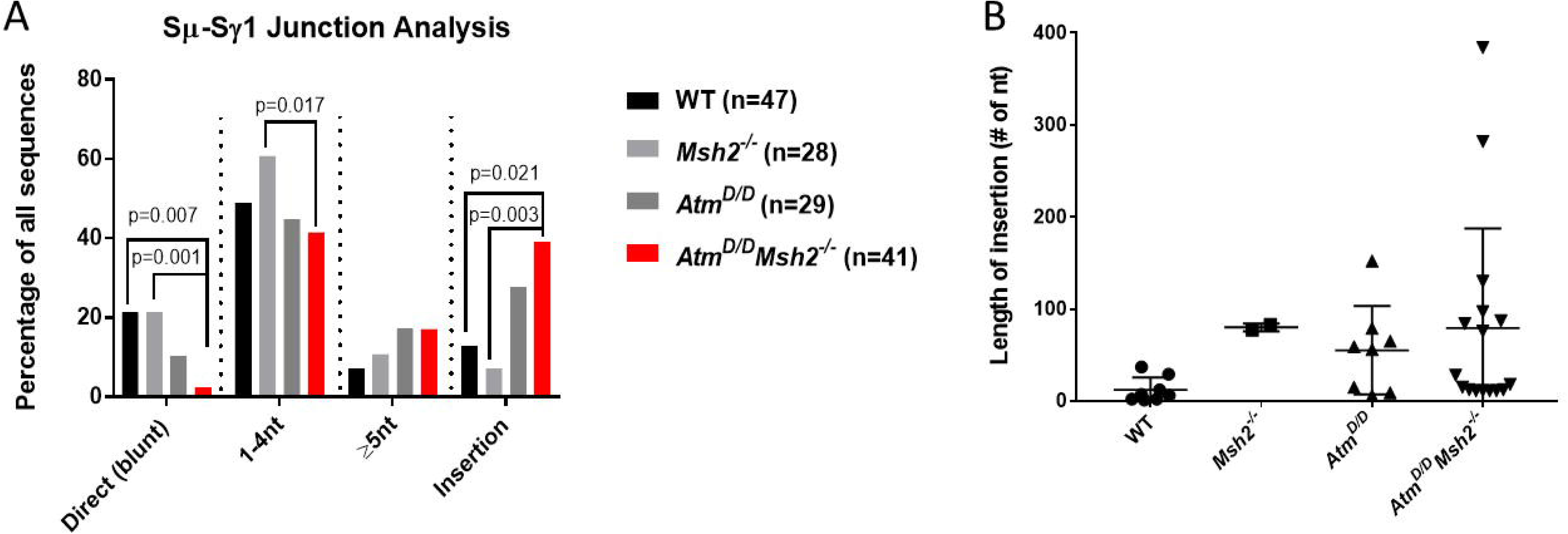
Significant loss of blunt end joins and increased insertions in *Atm^D/D^Msh2^-/-^*B cells undergoing CSR to IgG1. (**A**) Percentage of Sμ-Sγ1 junctions with blunt end join (0 nucleotide overlap), 1-4 nucleotide microhomology, ≥ 5 nucleotide microhomology, or insertion for the indicated genotype. Statistical analysis was conducted using a pairwise t-test with Holm-Sidak correction. Reported p-values are unadjusted. (**B**) Length of insertions in Sμ-Sγ1 junctions for the indicated genotype. Error bars mean (n ≥ 3 mice for each genotype). No statistical difference observed by one way ANOVA.

Significant differences in blunt end joins and insertions are also observed between *Atm^D/D^Msh2^-/-^*and *Msh2^-/-^* B cells (p=0.001 and p=0.003, respectively). Although we observed a smaller total percentage of insertions at Sμ-Sγ1 junctions in *Msh2^-/-^* B cells as compared to previous Sμ-Sγ3 analysis [10], these insertions were more than triple the length of that observed in WT Sμ-Sγ1 junctions (**Figure 3B**). The average inserted sequence at Sμ-Sγ1 junctions in *Msh2^-/-^* B cells was 80nt as compared to 12nt in WT B cells. Similarly, *Atm^D/D^* insertion lengths were longer (average 55nt) (**Figure 3B**). Interestingly, the Sμ-Sγ1 junctions in the *Atm^D/D^Msh2^-/-^*B cells skew towards insertion lengths of 100nt or longer, with the largest insertion size of almost 400nt; however, the average insertion lengths in Sμ-Sγ1 junctions of *Atm^D/D^Msh2^-/-^* B cells was not statistically different from the average insertion lengths in *Atm^D/D^*and *Msh2^-/-^* Sμ-Sγ1 junctions. The significant loss of blunt end joins coupled with an increase in insertions observed in Sμ-Sγ1 junctions of *Atm^D/D^Msh2^-/-^* B cells indicates that MSH2 enforces blunt end joining in the absence of ATM and identifies a distinct recombinogenic repair in *Atm^D/D^Msh2^-/-^* B cells as compared to *Atm^D/D^* or *Msh2^-/-^* B cells. Collectively, these data suggest that the DSBs in *Atm^D/D^Msh2^-/-^*S regions are joined in a process even more inefficient than either NHEJ or A-EJ that result in an inefficient and sloppy end joining.

## Discussion

Phosphorylation of AID and the subsequent interaction of pS38-AID with APE1 depends on ATM [19], suggesting that ATM may regulate CSR through BER. Based on genetic crosses that inactivated BER and MMR [8], genetic inactivation of *Atm* and *Msh2* was predicted to block both BER and MMR pathways, respectively, and abrogate CSR. Surprisingly, dihybrid crosses of *Atm^+/-^Msh2^+/-^* yielded no adult *Atm^-/-^Msh2^-/-^*mice, suggesting a synthetic lethal interaction between mutations in *Atm* and *Msh2* (**Table 1**). However, *Msh2^-/-^* mice with a conditional deletion of *Atm* in B cells (*Atm^D/D^Msh2^-/-^*) are viable in adulthood, indicating that the *Atm^-/-^Msh2^-/-^* lethality is not B cell dependent. Although the loss of *Atm* and *Msh2* induces a late embryonic to early postnatal lethal phenotype in mice (**Tables 2 and 3**), the underlying cellular mechanism behind this lethality remains unknown. Synthetic lethality (SL) results when the simultaneous loss of two genes leads to cell death, whereas the loss of each gene alone is tolerated and the cells are viable [50–52]. In mice, SLs between genes encoding DNA damage response proteins usually occur during gestation and early postnatal [53–57]. For example, mutations in the genes for DNA-PKcs and ATM lead to a synthetic lethal phenotype and death of mouse embryos at E7 [58]. Interestingly, loss of XRCC4 and DNA Ligase IV results in a late embryonic lethality similar to the *Atm^-/-^Msh2^-/-^* embryos [59, 60]. The significant loss of blunt end joins in *Atm^D/D^Msh2^-/-^* B cells suggests a block in NHEJ in the absence of ATM and MSH2 (**Figure 3A**). However, whether ATM and MSH2 deficient mouse embryos exhibit any defects in genomic instability and consequently cell proliferation, cell cycle arrest, or cell death remains to be determined. Alternatively, the reduction in observed *Atm^-/-^Msh2^-/-^*mice could result from metabolic defects. ATM and MSH2 respond directly or indirectly to reactive oxidative species (ROS), which are produced as byproducts of respiration [61, 62]. *Atm^-/-^* cells contain higher levels of intracellular ROS as compared to WT cells and the mechanism by which ATM responds to ROS is independent of its role in DSB [63, 64]. Additionally, MSH2 deficient cells have increased 8-oxo-G containing DNA, a byproduct of oxidative damage [65]. Thus, the synthetic lethality between *Atm* and *Msh2* may result from cellular respiratory failure or mitochondrial damage.

Contrary to the original hypothesis, *Atm^D/D^Msh2^-/-^*B cells complete CSR, albeit at severely impaired levels (**Figures 1-2**, **Supplemental Figure 5**). These data suggest ATM functions upstream of BER and MMR during CSR. Alternatively, ATM may be dispensable for BER-driven CSR. Interestingly, *Atm^-/-^Ung^-/-^*mice develop to adulthood and are obtained at Mendelian frequencies (data not shown). *Atm^-/-^Ung^-/-^* mouse IgG1 titers are comparable to *Atm^-/--^*mouse IgG1 titers (data not shown), further suggesting that ATM functions epistatically to BER and MMR. Contrary to the hypothesized role of ATM modulating BER through pS38-AID and APE1, IgG1 titers in *Atm^D/D^Msh2^-/-^* mice were significantly reduced pre- and post-immunization but not ablated (**Figures 1, 2, and Supplemental Figure 5**). To examine the DNA repair pathway utilized in the absence of ATM and MSH2, Sμ-Sγ1 junctions were sequenced in *Atm^D/D^Msh2^-/-^* B cells that were stimulated to switch to IgG1. Despite the smaller sample size from Sanger sequencing (**Figure 3**), the Sμ-Sγ1 junction data are comparable to a high-throughput genome wide sequencing of stimulated B cells [66] and confirms that ATM promotes NHEJ of S-S junctions in IgG1-switched B cells. In contrast to previous data published for S_μ_-S 3 junctions [10], *Msh2^-/-^* B cells show Sμ-Sγ1 junctions that are comparable to WT B cells (**Figure 3A**). However, in the absence of the NHEJ factor XRCC4, ablating MMR further decreases CSR, supporting the hypothesis that MMR can drive A-EJ [25]. Interestingly, the reduced CSR in B cells doubly deficient for XRCC4 and MSH2 is comparable to CSR in the *Atm^D/D^Msh2^-/-^* B cells (**Figure 2**, **Supplementary Figure 3**) [25]. Furthermore, blunt end joins in Sμ-Sγ1 junctions of *Atm^D/D^Msh2^-/-^* B cells are almost negligible, which indicates a block in NHEJ (**Figure 3A**) and is consistent with the phenotype observed in the NHEJ deficient *Xrcc4^-/-^*B cells [25]. Collectively, this would suggest MSH2 may enforce blunt end joining in the absence of NHEJ [25]. Speculatively, NHEJ and A-EJ may function sub-optimally in ATM and MSH2 deficient B cells to create a heterogeneous phenotype in end joining where the absence of ATM and MSH2 exert an epistatic suppression of NHEJ, which may account for the reduction in blunt joins. Alternatively, other NHEJ proteins may compensate for the absence of ATM [58, 67].

DNA-PKcs, a homolog of ATM, can partially function in the absence of ATM [68], which could account for the lack of significant difference in the percentage of short stretches of microhomologies (1-4nt) in *Atm^D/D^Msh2^-/-^*B cells as compared to the control genotypes (**Figure 3**). However, the extent to which DNA-PKcs can compensate for the absence of ATM may depend upon the amount of resection at the broken ends. When limited resection of DSB ends has occurred, the Ku70/80 complex efficiently binds the DSB [69], which allows for recruitment of DNA-PKcs, Artemis, XRCC4, and LigIV to efficiently join the broken ends through NHEJ [70, 71]. Because the Ku complex has low affinity for resected DNA [72], the initiation of resection is a critical determinant for repair pathway choice. Interestingly, there is evidence to suggest A-EJ driven by resection occurs in CSR. RPA, a ssDNA binding protein that regulates DNA resection [73, 74], interacts with pS38-AID at switch regions [20]. Furthermore, B cells defective in 53BP1 exhibit an increase in ATM-mediated end resection that is repaired through A-EJ during CSR [75]. Interestingly, *Atm^D/D^Msh2^-/-^* B cells show an increased percentage of Sμ- Sγ1 junctions with insertions, many of which are some of the longest in length (up to 380bp) across all the assayed genotypes (**Figure 3B**). This phenotype is comparable to that observed in the absence of 53BP1, suggesting that *Atm^D/D^Msh2^-/-^* insertions may have an increase in end resections that have not been repaired through A-EJ. An alternative explanation for the increased insertions could be that the absence of ATM and MSH2 leads to an increase in resection with no mechanism to repair through NHEJ or A-EJ and thus requires the need for a salvage pathway. Single strand annealing (SSA) has been observed in yeast in the case of resected DNA homology-based repair but in the absence of functional HR, leading to high rates of mutagenesis [76–78]. In this way, the increase in insertions observed in *Atm^D/D^Msh2^-/-^* B cells may indicate an attempt at salvage repair through SSA. Collectively, this may indicate that ATM and MSH2 are required to repair resected DNA to drive A-EJ (**Figure 4**).

**Figure 4:**
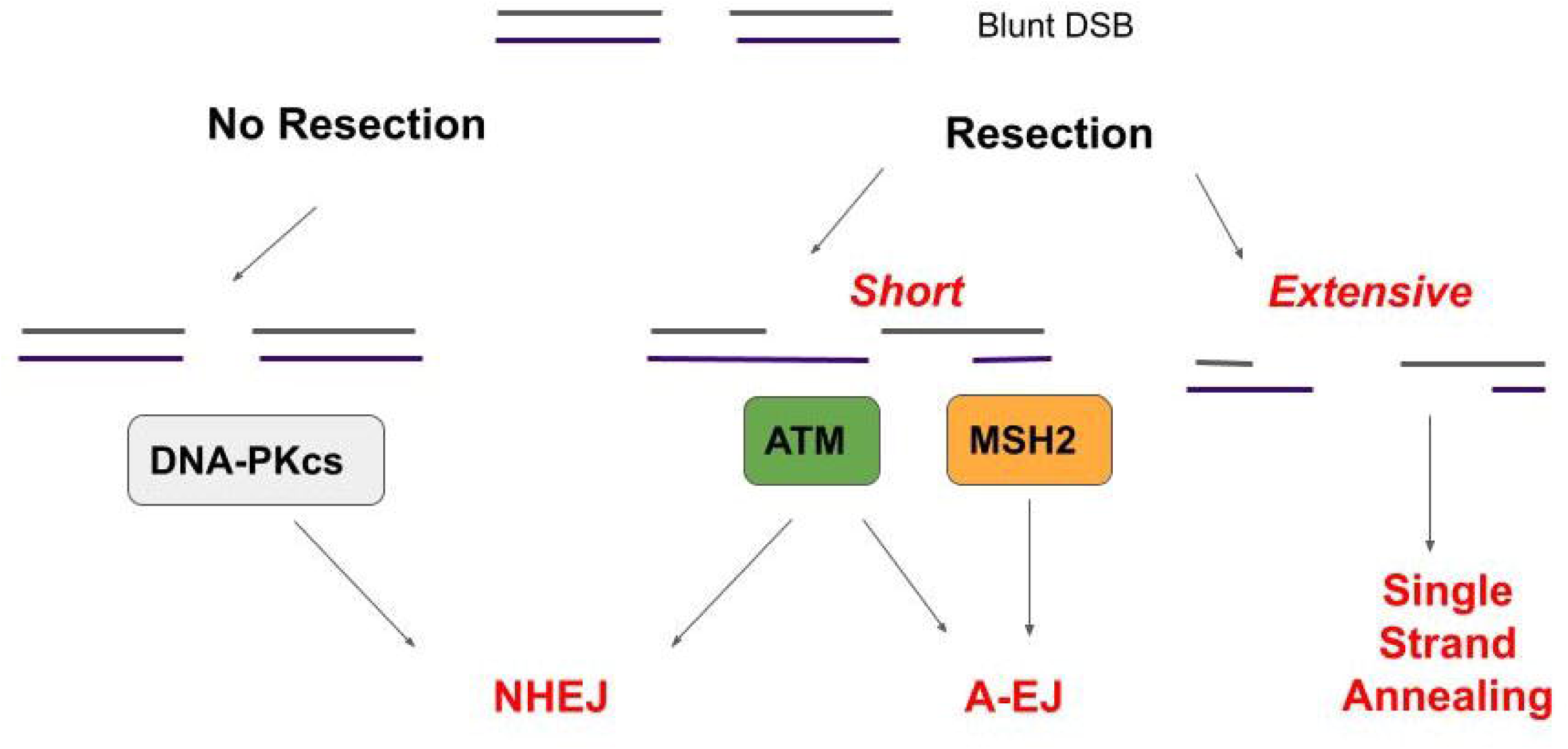
A model for the requirement of ATM and MSH2 to repair resected DNA. In NHEJ, blunt end joins are repaired through DNA-PKcs. However, when DSBs are resected, NHEJ repair is mediated by ATM, which could direct phosphorylation of AID at S38 to recruit RPA-coated ssDNA and promote MMR to carry out microhomology-mediated A-EJ. In the absence of ATM and MSH2, resected ends may be repaired through a salvage pathway like single strand annealing (SSA) in an attempt at homology-driven repair.

We hypothesize that pS38-AID mediated interaction with RPA-coated resected ssDNA may signal through ATM in a positive feedback loop to increase the density of resected DNA breaks in recombining switch regions [19, 79]. We propose a model whereby the phosphorylation of AID through an uncharacterized ATM-PKA signaling cascade activates A-EJ in CSR (**Figure 4**). Hypothetically, in the absence of resection (i.e. blunt end joins), the Ku complex efficiently binds the broken ends and recruits DNA-PKcs mediated NHEJ. However, when the DSB ends have been resected, NHEJ is driven by ATM. We speculate that activation of ATM indirectly induces the phosphorylation of S38 on AID, which in turn may co-interact with resected SSB coated by RPA [80], facilitating microhomology-mediated strand invasion into the acceptor switch region to complete repair through A-EJ. We speculate that in the absence of ATM and MSH2, resected DNA can only be repaired through SSA.

## Supplemental Figure Legends

**Supplemental Table 1:** The numbers of expected and observed progeny from *CD23cre^+^Atm^f/f^Msh2^+/-^* X *Atm^f/f^Msh2^+/-^* or *CD23cre^+^Atm^f/f^Msh2^+/-^* X *Atm^f/f^Msh2^-/-^* at P28. P-value determined by χ^2^ test.

**Supplemental Figure 1: *Atm^D/D^Msh2^-/-^* mice show no significant alteration in germinal center (GC) B cell numbers**. Peyer’s patch cells were harvested 10 days after NP-CCG immunization and stained with anti-B220 antibody and peanut agglutinin (PNA). Percentage of germinal center B cells (B220^+^PNA^+^) from each mouse of the indicated genotype is plotted. Statistical analysis was completed by one way ANOVA (*= p<0.05) and error bars indicate the mean and standard deviation. n ≥ 3 mice per genotype.

**Supplemental Figure 2: Immunoblot of protein extracts from LPS plus IL4-stimulated B cells shows significant loss of ATM and MSH2 in *Atm^D/D^Msh2^-/-^* B cells.** Protein extracts were prepared from B cells at 72 hours post-stimulation, resolved on either an 8% SDS-PAGE (ATM, MSH2, αtubulin) or 10% SDS-PAGE (AID, αtubulin), and transferred to PVDF. Immunoblots were probed with anti-ATM, anti-MSH2, anti-AID, and anti-αtubulin antibodies.

**Supplemental Figure 3: *Atm^D/D^Msh2^-/-^*B cells and mice show reduced IgG3 CSR.** (**A**) LPS stimulated B cells of the indicated genotypes were stained with anti-IgG3 antibody and assayed by flow cytometry. The percentage of IgG3^+^ cells is plotted for each genotype. (**B**) Serum titers of IgG3 were assayed before or after immunization with NP-CGG. Statistical analysis was conducted by one way ANOVA (* p<0.05, ** p<0.005, ** p<0.005, ***p<0.0005, ****p<.00001). The error bars indicate mean and standard deviation. n ≥ 3 mice per genotype.

**Supplemental Figure 4: No significant alteration in *Atm^D/D^Msh2^-/-^*B cell proliferation**. LPS+IL4 stimulated B cells were labeled with SNARF. Cells were analyzed via flow cytometry every 24 hours for two days. Mean fluorescence intensity (MFI) of SNARF fluorescence in the PE-TX Red channel, was quantified for each time point and normalized to the D0 WT control. Statistical analysis was conducted by student’s t-test (**p<0.0005). The error bars indicate mean and standard deviation. n ≥ 3 mice per genotype.

**Supplemental Figure 5: *Atm^D/D^Msh2^-/-^* B cells display no significant alterations in germline transcription (GLT)**. (**A**) Aggregate fold expression of μ GLT from LPS+IL4 stimulated B cells normalized to expression of βactin and WT B cells. (**B**) Aggregate fold expression of γ1 GLT from LPS+IL4 stimulated B cells normalized to expression of βactin and WT B cells. n ≥ 3 mice per genotype.

## Materials and Methods

### Mice

C57BL/6 and *Atm^-/-^* (stock #008536) mice were purchased from The Jackson Laboratory [34, 81]. *Msh2^-/-^*were a gift from H. te Riele [35]. *Aicda^-/-^* mice were a gift from T. Honjo [42]. *Atm^fl/f^*mice were a gift of Shan Zha [39]; and *CD23-Cre* mice were a gift of Meinrad Busslinger [82]. Husbandry of and experiments with mice were conducted according to protocols approved by The City College of New York Institutional Animal Care and Use Committee.

### Genotyping of mice and embryos

To obtain E13.5 embryos, females 3-6 months of age were placed into a cage previously occupied by male mice one day prior to introducing the male mouse. Timed matings were set up by limiting the mating period to one day. Pregnant female mice were euthanized 14 days after mating (E13.5) and embryos were collected from the uterine horn immediately after euthanasia. To genotype the embryos, one hind limb and tail were incubated in lysis buffer (10 mM Tris, 0.1 M EDTA, 0.5% SDS, and 200μg/ml of Proteinase K) overnight at 56°C. Genomic DNA was isolated by isopropanol precipitation and resuspended in TE buffer. To genotype newborn mice (P0), pregnant females were monitored daily for neonates approximately three weeks after mating. Tail biopsies were collected from neonates for genotyping. If a dead carcass or still born neonate was found in the litter, tissue samples or tail biopsies were collected and genotyped when possible. Adult mice were identified by tail biopsy approximately four weeks after birth (P28), when the mice were weaned from the nursing female. The primers for each genotype are as follows: *Atm*[ATMB 5’-CGAATTTGCAGGAGTTGCTGAG-3’, ATMF:5’ GACTTCTGTCAGATGTTGCTGCC-3’, and ATMN:5’-

GGGTGGGATTAGATAAATGCCTG-3’], *Msh2* [MshF: 5’- GCTCACTTAGACGCCATTGT- 3’, MshWR: 5’-AAAGTGCACGTCATTTGGA-3’, and MshKR: 5’- GCCTTCTTGACGAGTTCTTC-3’], CD23-Cre [CD23cre_1: 5’- GCGACTAGTAACTTGTTTATTGCA GCTTAT-3’, CD23cre_2: 5’- GCGTCCGGAAGACACCACATCCCAATTCTT -3’], and *Atm^f^* [ATMgF86723: 5’- ATCAAATGTAAAGGCGGCTTC-3’, ATM_BAC13: 5’- CATCCTTTAATGTGCCTCCCTTCGCC-3’, ATM_BAC7: 5’-GCCCATCCCGTCCACAATATCTCTGC-3’]. PCR with DreamTaq (Fisher #K1082) was performed for 30 cycles using the following conditions: 95°C, 30sec; Tm, 30 sec; 72°C, 30sec. The Tm (annealing temperature) for *Atm* and *Msh2* was 55°C, CD23-Cre 58°C, and *Atm^f^* 63°C.

### B cell purification and stimulation for CSR

B cells were purified from the spleens of mice (aged 2–7 months of age) through negative selection with anti-CD43 magnetic beads (Miltenyi Biotec) and subsequently cultured in RPMI 1640 media (Life Technologies) supplemented with 10% FBS (Atlanta Biologicals), 2 mM L-glutamine, 1x penicillin/streptomycin (Corning), and 47.3 μM 2-mercaptoethanol. For switching to IgG1, B cells cultured at 1x10^6^ cells/ml with 25 μg/ml LPS (catalog no. L7261; Sigma-Aldrich) and 12.5 ng/ml IL-4 (R&D Systems). B cells were split 1:2 v/v 48 and 72 hours after isolation. To stimulate CSR to IgG3, 25 μg/ml LPS was used. After 96 hours post-stimulation, B cells were harvested for analysis by flow cytometry. Live cells were kept on ice and stained in Gibco Phosphate Buffered Saline (PBS) pH 7.4 (catalog no. 10-010-049) supplemented with 2.5% FBS. To identify switched B cells, samples were stained with anti-IgG1 (BD; APC conjugate catalog no. BDB550874 or PE conjugate catalog no. BDB550083) or anti-IgG3 (BD; FITC conjugate catalog no. BDB553403). DAPI was used for live dead exclusion. Samples were analyzed on a BD LSR II cytometer. For proliferation assays, B cells were labeled with Carboxylic Acid, Acetate, Succinimidyl Ester (SNARF) (ThermoFisher catalog no. S22801) and analyzed by flow cytometry at time of labeling (D0), and every 24 hours thereafter.

### Immunoblots and RNA expression

Protein extracts were prepared from IgG1 stimulated B cells at 72 hours post-stimulation. Cells were lysed in NP40 lysis buffer supplemented with proteinase inhibitor tablet (Millipore Sigma). NP40 cell lysates were resolved on 8% SDS PAGE gels, transferred overnight to Immobilon (Millipore Sigma), and immunoblotted with anti-ATM (GeneTex catalog no. GTX70103), anti-MSH2 (Abcam catalog no. ab70270), or anti-αtubulin (Sigma catalog no. T9026-100UL) antibodies. Immunoblots were developed using anti-mouse-HRP or anti-Rabbit-HRP using the Pierce ECL Western blotting substrate (Fisher catalog no. PI32106) and imaged on an Azure A300 chemiluminescent imaging system.

RNA was extracted from IgG1 stimulated B cells 72 hour post-stimulation using TRIzol reagent (Fisher) and cDNA was prepared using the Protoscript II kit (NEB). Germline transcription was analyzed using quantitative PCR with SYBR Green (Roche) and primers for Sμ (ImUF: 5’- CTCTGGCCCTGCTTATTGTTG-3’, CmuR: 5’-GAAGACATTTGGGAAGGACTGACT-3’) or Sγ1 (Igamma1: 5’- GGCCCTTCCAGATCTTTGAG-3’, Cgamma1: 5’-GGATCCAGAGTTCCAGGTCACT-3’). Reactions were analyzed on a Lightcycler96 (Roche).

### Mouse immunizations and Ig titers

Mice (aged 2–6 months) were injected intraperitoneally with 50 μg of 4-hydroxy-3-nitrophenylacetyl conjugated to chicken gamma globulin (NP-CGG) (Biosearch Technologies) in Imject Alum (Thermo Fisher Scientific) [83]. Mice were given a boost of NP-CGG in Imject Alum 10 days post–primary immunization. Sera were obtained from blood that was collected on day 0 (pre-immunization) and day 7 and day 21 (post-immunization) through cheek bleeds (days 0 and 7) or cardiac puncture (day 21). To analyze Ig serum titers, ELISA assays were performed as previously described [22].

### Switch Junction Analysis

DNA from IgG1-stimulated B cells was isolated from ∼1.5x10^6^ cells 72hours after stimulation. Sμ-Sγ1 junctions were amplified using nested PCR primers: Smu1b (5’- AACTCTCCAGCCACAGTAATGACC-3’) and Sγ1.1 (5’-CTGTAACCTACCCAGGAGACC-3’) were used for the first cycle of PCR; SmuF2 (5’-GAGAAGGCCAGACTCATAAAGCT-3’) and Sγ1.1R2 (5’-GTCGAATTCCCCCATCCTGTCACCTATA-3’) were used in the second round PCR, as previously described [45, 84]. Both rounds of PCR were performed for an initial 10 cycles at 94°C for 15s, 52°C for 30s, 72°C 2 min followed by 23 more cycles with 5s added per cycle. All PCRs were performed using Q5 High Fidelity DNA Polymerase (NEB) according to the manufacturer’s instructions. Five identical PCR reactions were setup for each B cell stimulation. The reactions were pooled, concentrated in a speed-vac, resolved on a 1% agarose gel. PCR products (0.5-1kb) were excised from the gel, cloned into cloning vector pJET1.2 (Fisher catalog no. K1231), and Sanger sequenced using the T7 forward primer. Sequenced junctions were aligned against reference Sμ (MUSIGCD07) and Sγ1 (MUSIGHANB) in NCBI Blast with the low complexity filter disabled.

### Figure composition and Statistical analysis

Data figures were generated using Graphpad Prism 7.01. Unless otherwise noted, statistical analyses were completed by one way ANOVA in Prism. Graphical models were created through Biorender.

## Acknowledgements

This work was supported by funding from the National Cancer Institute (2U54CA132378) and National Institute of General Medical Sciences (1SC1GM132035). Mouse husbandry and colony maintenance was supported by Harry Acosta, Alfredo Diaz, Anthony Pacheco, and Iris Falquez from the CCNY Marshak Animal Care Facility. We thank Jayanta Chaudhuri for the anti-AID antibody. Thank you to Drs. Alicia Meléndez, Christine Li, Scott Keeney, Linda Spatz for providing intellectual support, and Dr. Allysia Matthews, Dr. Simin Zheng, and Sarah (Jee Eun) Choi for critical reading of the manuscript.

## Abbreviations

AID: activation induced cytidine deaminase
A-EJ: alternative end joining
APE1: apurinic/apyrimadinic endonuclease 1
ATM: ataxia telangiectasia mutated
BER: base excision repair pathway
CSR: Class switch recombination
DNA-PKcs: DNA-dependent protein kinase catalytic subunit
DSB: double stranded DNA break
E13.5: mouse embryonic day 13.5
MMR: mismatch repair pathway
pS38-AID: phosphorylation of serine 38 on AID
MSH2: mut S homolog 2
MLH1: mutL homology 1
NHEJ: Nonhomologous end joining
P0: newborn mouse pup
P28: Post natal day 28
PMS2: PMS1 homolog 2
SSA: single strand annealing
SSB: single stranded DNA break
UNG: uracil DNA glycosylase 1
XRCC4: X-ray repair cross complementing

## References

1. Alt, F.W., et al., VDJ recombination. Immunology today, 1992. 13(8): p. 306–314.

2. Chaudhuri, J. and F.W. Alt, Class-switch recombination: interplay of transcription, DNA deamination and DNA repair. Nature Reviews Immunology, 2004. 4(7): p. 541–552.

3. Rush, J.S., S.D. Fugmann, and D.G. Schatz, Staggered AID-dependent DNA double strand breaks are the predominant DNA lesions targeted to Sµ in Ig class switch recombination. International immunology, 2004. 16(4): p. 549–557.

4. Alt, F.W., et al., Mechanisms of programmed DNA lesions and genomic instability in the immune system. Cell, 2013. 152(3): p. 417–429.

5. Xu, Z., et al., Immunoglobulin class-switch DNA recombination: induction, targeting and beyond. Nature Reviews Immunology, 2012. 12(7): p. 517–531.

6. Methot, S. and J. Di Noia, Molecular mechanisms of somatic hypermutation and class switch recombination, in Advances in immunology. 2017, Elsevier. p. 37-87.

7. Honjo, T., M. Muramatsu, and S. Fagarasan, AID: how does it aid antibody diversity? Immunity, 2004. 20(6): p. 659–668.

8. Rada, C., J.M. Di Noia, and M.S. Neuberger, Mismatch recognition and uracil excision provide complementary paths to both Ig switching and the A/T-focused phase of somatic mutation. Molecular cell, 2004. 16(2): p. 163–171.

9. Rada, C., et al., Immunoglobulin isotype switching is inhibited and somatic hypermutation perturbed in UNG-deficient mice. Current Biology, 2002. 12(20): p. 1748–1755.

10. Schrader, C.E., J. Vardo, and J. Stavnezer, Role for mismatch repair proteins Msh2, Mlh1, and Pms2 in immunoglobulin class switching shown by sequence analysis of recombination junctions. The Journal of experimental medicine, 2002. 195(3): p. 367–373.

11. Martin, A., et al., Msh2 ATPase activity is essential for somatic hypermutation at AT basepairs and for efficient class switch recombination. The Journal of experimental medicine, 2003. 198(8): p. 1171–1178.

12. Imai, K., et al., Human uracil–DNA glycosylase deficiency associated with profoundly impaired immunoglobulin class-switch recombination. Nature immunology, 2003. 4(10): p. 1023–1028.

13. Masani, S., L. Han, and K. Yu, Apurinic/apyrimidinic endonuclease 1 is the essential nuclease during immunoglobulin class switch recombination. Molecular and cellular biology, 2013. 33(7): p. 1468–1473.

14. Guikema, J.E., et al., APE1-and APE2-dependent DNA breaks in immunoglobulin class switch recombination. The Journal of experimental medicine, 2007. 204(12): p. 3017–3026.

15. Schrader, C.E., et al., The roles of APE1, APE2, DNA polymerase β and mismatch repair in creating S region DNA breaks during antibody class switch. Philosophical Transactions of the Royal Society B: Biological Sciences, 2009. 364(1517): p. 645–652.

16. Schrader, C.E., J. Vardo, and J. Stavnezer, Role for mismatch repair proteins Msh2, Mlh1, and Pms2 in immunoglobulin class switching shown by sequence analysis of recombination junctions. Journal of Experimental Medicine, 2002. 195(3): p. 367–373.

17. Seeberg, E., L. Eide, and M. Bjørås, The base excision repair pathway. Trends in biochemical sciences, 1995. 20(10): p. 391–397.

18. Tishkoff, D.X., et al., Identification and characterization of Saccharomyces cerevisiae EXO1, a gene encoding an exonuclease that interacts with MSH2. Proceedings of the National Academy of Sciences, 1997. 94(14): p. 7487–7492.

19. Vuong, B.Q., et al., A DNA break–and phosphorylation-dependent positive feedback loop promotes immunoglobulin class-switch recombination. Nature immunology, 2013. 14(11): p. 1183–1189.

20. Cheng, H.-L., et al., Integrity of the AID serine-38 phosphorylation site is critical for class switch recombination and somatic hypermutation in mice. Proceedings of the National Academy of Sciences, 2009. 106(8): p. 2717–2722.

21. McBride, K.M., et al., Regulation of class switch recombination and somatic mutation by AID phosphorylation. The Journal of experimental medicine, 2008. 205(11): p. 2585–2594.

22. Choi, J.E., et al., AID Phosphorylation Regulates Mismatch Repair–Dependent Class Switch Recombination and Affinity Maturation. The Journal of Immunology, 2020. 204(1): p. 13–22.

23. Reina-San-Martin, B., et al., ATM is required for efficient recombination between immunoglobulin switch regions. The Journal of experimental medicine, 2004. 200(9): p. 1103–1110.

24. Lumsden, J.M., et al., Immunoglobulin class switch recombination is impaired in Atm-deficient mice. Journal of Experimental Medicine, 2004. 200(9): p. 1111–1121.

25. Eccleston, J., et al., Mismatch repair proteins MSH2, MLH1, and EXO1 are important for class-switch recombination events occurring in B cells that lack nonhomologous end joining. The Journal of Immunology, 2011. 186(4): p. 2336–2343.

26. Rooney, S., J. Chaudhuri, and F.W. Alt, The role of the non-homologous end-joining pathway in lymphocyte development. Immunological reviews, 2004. 200(1): p. 115–131.

27. Boboila, C., et al., Alternative end-joining catalyzes class switch recombination in the absence of both Ku70 and DNA ligase 4. Journal of Experimental Medicine, 2010. 207(2): p. 417–427.

28. Yan, C.T., et al., IgH class switching and translocations use a robust non-classical end-joining pathway. Nature, 2007. 449(7161): p. 478–482.

29. Soulas-Sprauel, P., et al., Role for DNA repair factor XRCC4 in immunoglobulin class switch recombination. Journal of Experimental Medicine, 2007. 204(7): p. 1717–1727.

30. Boboila, C., et al., Robust chromosomal DNA repair via alternative end-joining in the absence of X-ray repair cross-complementing protein 1 (XRCC1). Proceedings of the National Academy of Sciences, 2012. 109(7): p. 2473–2478.

31. Boboila, C., et al., Alternative end-joining catalyzes robust IgH locus deletions and translocations in the combined absence of ligase 4 and Ku70. Proceedings of the National Academy of Sciences, 2010. 107(7): p. 3034–3039.

32. Dunnick, W., et al., DNA sequences at immunoglobulin switch region recombination sites. Nucleic Acids Research, 1993. 21(3): p. 365–372.

33. Pan, Q., et al., Alternative end joining during switch recombination in patients with Ataxia-Telangiectasia. European journal of immunology, 2002. 32(5): p. 1300–1308.

34. Barlow, C., et al., Atm-deficient mice: a paradigm of ataxia telangiectasia. Cell, 1996. 86(1): p. 159–171.

35. de Wind, N., et al., Inactivation of the mouse Msh2 gene results in mismatch repair deficiency, methylation tolerance, hyperrecombination, and predisposition to cancer. Cell, 1995. 82(2): p. 321–330.

36. Reitmair, A., et al., MSH2 deficient mice are viable and susceptible to lymphoid tumours. Nature genetics, 1995. 11(1): p. 64–70.

37. Callén, E., et al., ATM prevents the persistence and propagation of chromosome breaks in lymphocytes. Cell, 2007. 130(1): p. 63–75.

38. Aubry, J.-P., et al., CD21 is a ligand for CD23 and regulates IgE production. Nature, 1992. 358(6386): p. 505–507.

39. Zha, S., et al., Complementary functions of ATM and H2AX in development and suppression of genomic instability. Proceedings of the National Academy of Sciences, 2008. 105(27): p. 9302–9306.

40. Yang Shih, T.-A., et al., Role of BCR affinity in T cell–dependent antibody responses in vivo. Nature immunology, 2002. 3(6): p. 570–575.

41. Toellner, K.-M., et al., Low-level hypermutation in T cell–independent germinal centers compared with high mutation rates associated with T cell–dependent germinal centers. The Journal of experimental medicine, 2002. 195(3): p. 383–389.

42. Muramatsu, M., et al., Class switch recombination and hypermutation require activation-induced cytidine deaminase (AID), a potential RNA editing enzyme. Cell, 2000. 102(5): p. 553–563.

43. Manis, J.P., M. Tian, and F.W. Alt, Mechanism and control of class-switch recombination. Trends in immunology, 2002. 23(1): p. 31–39.

44. Lee-Theilen, M., et al., CtIP promotes microhomology-mediated alternative end joining during class-switch recombination. Nature structural & molecular biology, 2011. 18(1): p. 75–79.

45. Ehrenstein, M.R., et al., Switch junction sequences in PMS2-deficient mice reveal a microhomology-mediated mechanism of Ig class switch recombination. Proceedings of the National Academy of Sciences, 2001. 98(25): p. 14553–14558.

46. Stavnezer, J., et al., Mapping of switch recombination junctions, a tool for studying DNA repair pathways during immunoglobulin class switching. Advances in immunology, 2010. 108: p. 45–109.

47. Bothmer, A., et al., 53BP1 regulates DNA resection and the choice between classical and alternative end joining during class switch recombination. Journal of Experimental Medicine, 2010. 207(4): p. 855–865.

48. Symington, L.S. and J. Gautier, Double-strand break end resection and repair pathway choice. Annual review of genetics, 2011. 45: p. 247–271.

49. Ehrenstein, M.R. and M.S. Neuberger, Deficiency in Msh2 affects the efficiency and local sequence specificity of immunoglobulin class-switch recombination: parallels with somatic hypermutation. The EMBO journal, 1999. 18(12): p. 3484–3490.

50. Ashworth, A., C.J. Lord, and J.S. Reis-Filho, Genetic interactions in cancer progression and treatment. Cell, 2011. 145(1): p. 30–38.

51. Farmer, H., et al., Targeting the DNA repair defect in BRCA mutant cells as a therapeutic strategy. Nature, 2005. 434(7035): p. 917–921.

52. Bryant, H.E., et al., Specific killing of BRCA2-deficient tumours with inhibitors of poly (ADP-ribose) polymerase. Nature, 2005. 434(7035): p. 913–917.

53. Kantidze, O.L., et al., Synthetically lethal interactions of ATM, ATR, and DNA-PKcs. Trends in cancer, 2018. 4(11): p. 755–768.

54. Daniel, J.A., et al., Loss of ATM kinase activity leads to embryonic lethality in mice. Journal of Cell Biology, 2012. 198(3): p. 295–304.

55. Murcia, J.M.n.-d., et al., Early embryonic lethality in PARP-1 Atm double-mutant mice suggests a functional synergy in cell proliferation during development. Molecular and cellular biology, 2001. 21(5): p. 1828–1832.

56. Huber, A., et al., PARP-1, PARP-2 and ATM in the DNA damage response: functional synergy in mouse development. DNA repair, 2004. 3(8-9): p. 1103–1108.

57. Sekiguchi, J., et al., Genetic interactions between ATM and the nonhomologous end-joining factors in genomic stability and development. Proceedings of the National Academy of Sciences, 2001. 98(6): p. 3243–3248.

58. Gurley, K.E. and C.J. Kemp, Synthetic lethality between mutation in Atm and DNA-PKcs during murine embryogenesis. Current Biology, 2001. 11(3): p. 191–194.

59. Gao, Y., et al., A critical role for DNA end-joining proteins in both lymphogenesis and neurogenesis. Cell, 1998. 95(7): p. 891–902.

60. Frank, K.M., et al., DNA ligase IV deficiency in mice leads to defective neurogenesis and embryonic lethality via the p53 pathway. Molecular cell, 2000. 5(6): p. 993–1002.

61. Kozlov, S.V., et al., Reactive oxygen species (ROS)-activated ATM-dependent phosphorylation of cytoplasmic substrates identified by large-scale phosphoproteomics screen. Molecular & Cellular Proteomics, 2016. 15(3): p. 1032–1047.

62. Martin, S.A., et al., DNA polymerases as potential therapeutic targets for cancers deficient in the DNA mismatch repair proteins MSH2 or MLH1. Cancer cell, 2010. 17(3): p. 235-248.

63. Ito, K., et al., Regulation of reactive oxygen species by Atm is essential for proper response to DNA double-strand breaks in lymphocytes. The Journal of Immunology, 2007. 178(1): p. 103–110.

64. Guo, Z., et al., ATM activation by oxidative stress. Science, 2010. 330(6003): p. 517-521.

65. Colussi, C., et al., The mammalian mismatch repair pathway removes DNA 8-oxodGMP incorporated from the oxidized dNTP pool. Current biology, 2002. 12(11): p. 912–918.

66. Panchakshari, R.A., et al., DNA double-strand break response factors influence end-joining features of IgH class switch and general translocation junctions. Proceedings of the National Academy of Sciences, 2018. 115(4): p. 762–767.

67. Callén, E., et al., Essential role for DNA-PKcs in DNA double-strand break repair and apoptosis in ATM-deficient lymphocytes. Molecular cell, 2009. 34(3): p. 285–297.

68. Cook, A.J., et al., Reduced switching in SCID B cells is associated with altered somatic mutation of recombined S regions. The Journal of Immunology, 2003. 171(12): p. 6556–6564.

69. Ira, G., et al., DNA end resection, homologous recombination and DNA damage checkpoint activation require CDK1. Nature, 2004. 431(7011): p. 1011–1017.

70. Blackford, A.N. and S.P. Jackson, ATM, ATR, and DNA-PK: the trinity at the heart of the DNA damage response. Molecular cell, 2017. 66(6): p. 801–817.

71. Falck, J., J. Coates, and S.P. Jackson, Conserved modes of recruitment of ATM, ATR and DNA-PKcs to sites of DNA damage. Nature, 2005. 434(7033): p. 605–611.

72. Dynan, W.S. and S. Yoo, Interaction of Ku protein and DNA-dependent protein kinase catalytic subunit with nucleic acids. Nucleic acids research, 1998. 26(7): p. 1551–1559.

73. Chen, H., M. Lisby, and L.S. Symington, RPA coordinates DNA end resection and prevents formation of DNA hairpins. Molecular cell, 2013. 50(4): p. 589–600.

74. Soniat, M.M., et al., RPA phosphorylation inhibits DNA resection. Molecular cell, 2019. 75(1): p. 145–153. e5.

75. Reina-San-Martin, B., et al., Enhanced intra-switch region recombination during immunoglobulin class switch recombination in 53BP1–/–B cells. European journal of immunology, 2007. 37(1): p. 235–239.

76. Weinstock, D.M., et al., Modeling oncogenic translocations: distinct roles for double-strand break repair pathways in translocation formation in mammalian cells. DNA repair, 2006. 5(9-10): p. 1065–1074.

77. Moynahan, M.E. and M. Jasin, Mitotic homologous recombination maintains genomic stability and suppresses tumorigenesis. Nature reviews Molecular cell biology, 2010. 11(3): p. 196–207.

78. Spies, M. and R. Fishel, Mismatch repair during homologous and homeologous recombination. Cold Spring Harbor perspectives in biology, 2015. 7(3): p. a022657.

79. Matthews, A.J., et al., Regulation of immunoglobulin class-switch recombination: choreography of noncoding transcription, targeted DNA deamination, and long-range DNA repair. Advances in immunology, 2014. 122: p. 1–57.

80. Yamane, A., et al., RPA accumulation during class switch recombination represents 5′–3′ DNA-end resection during the S–G2/M phase of the cell cycle. Cell reports, 2013. 3(1): p. 138–147.

81. Blunt, T., et al., Defective DNA-dependent protein kinase activity is linked to V (D) J recombination and DNA repair defects associated with the murine scid mutation. Cell, 1995. 80(5): p. 813–823.

82. Kwon, K., et al., Instructive role of the transcription factor E2A in early B lymphopoiesis and germinal center B cell development. Immunity, 2008. 28(6): p. 751–762.

83. Pucella, J.N., et al., miR-182 is largely dispensable for adaptive immunity: lack of correlation between expression and function. The Journal of Immunology, 2015. 194(6): p. 2635–2642.

84. Zan, H., et al., Rad52 competes with Ku70/Ku86 for binding to S-region DSB ends to modulate antibody class-switch DNA recombination. Nature communications, 2017. 8(1): p. 1–16.

