## Supplementary figures and images for "ATM and MSH2 control blunt DNA end joining in immunoglobulin class switch recombination"

### Supplemental figure 1

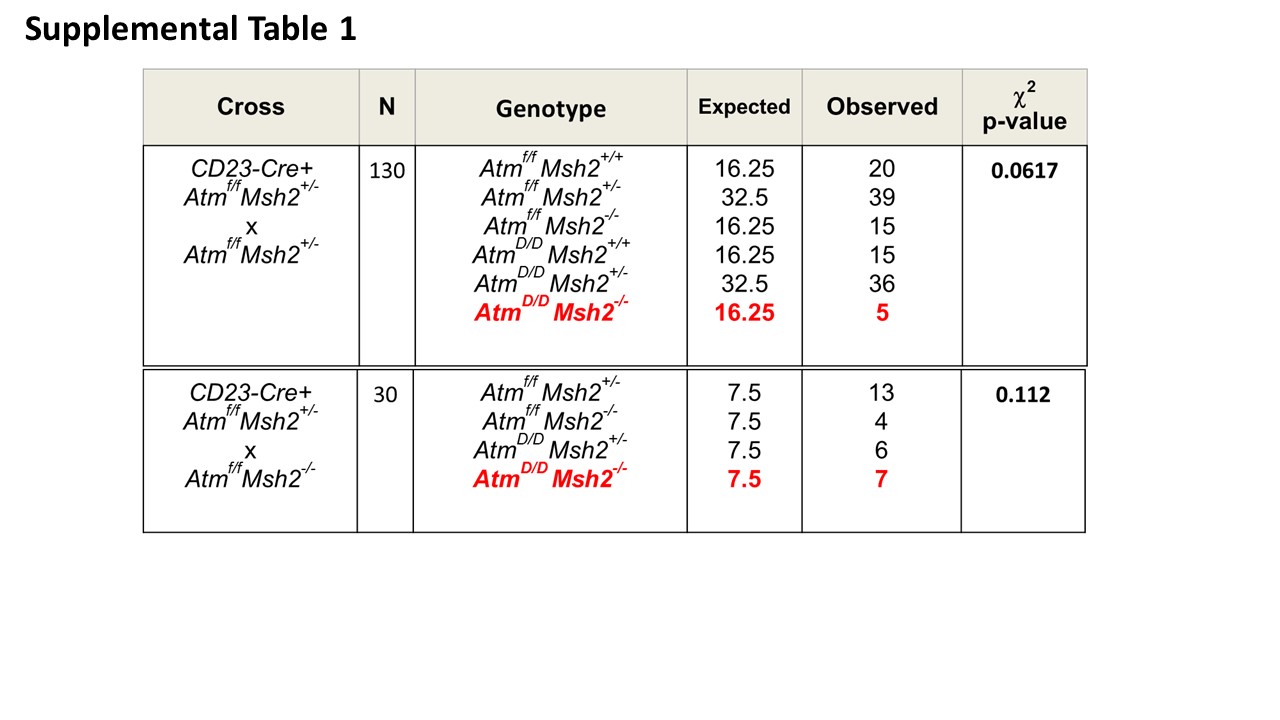

### Supplemental figure 2

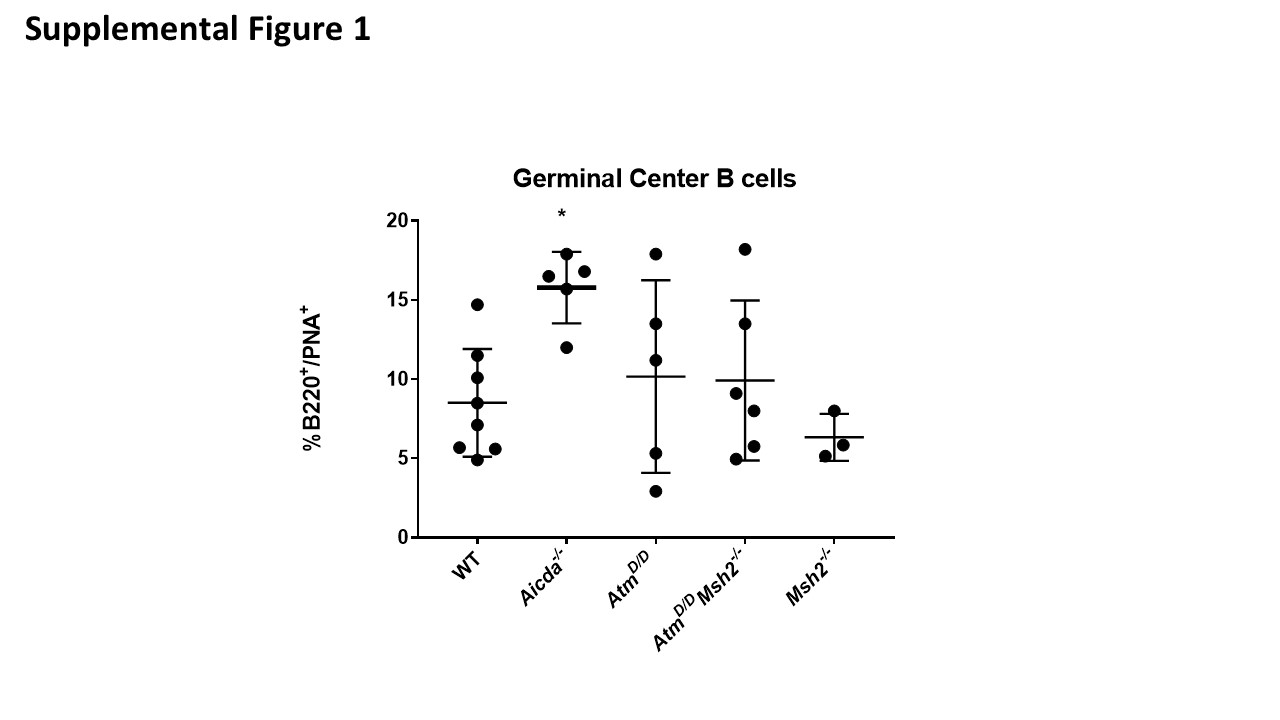

### Supplemental figure 3

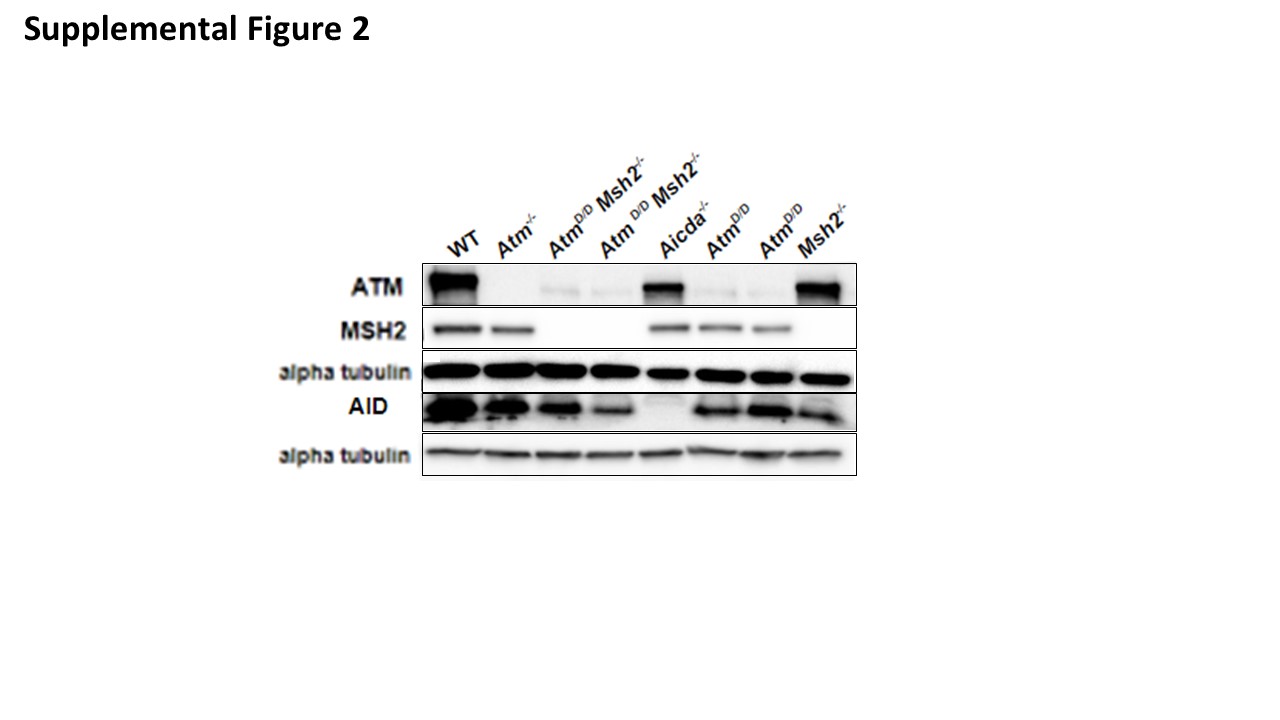

### Supplemental figure 4

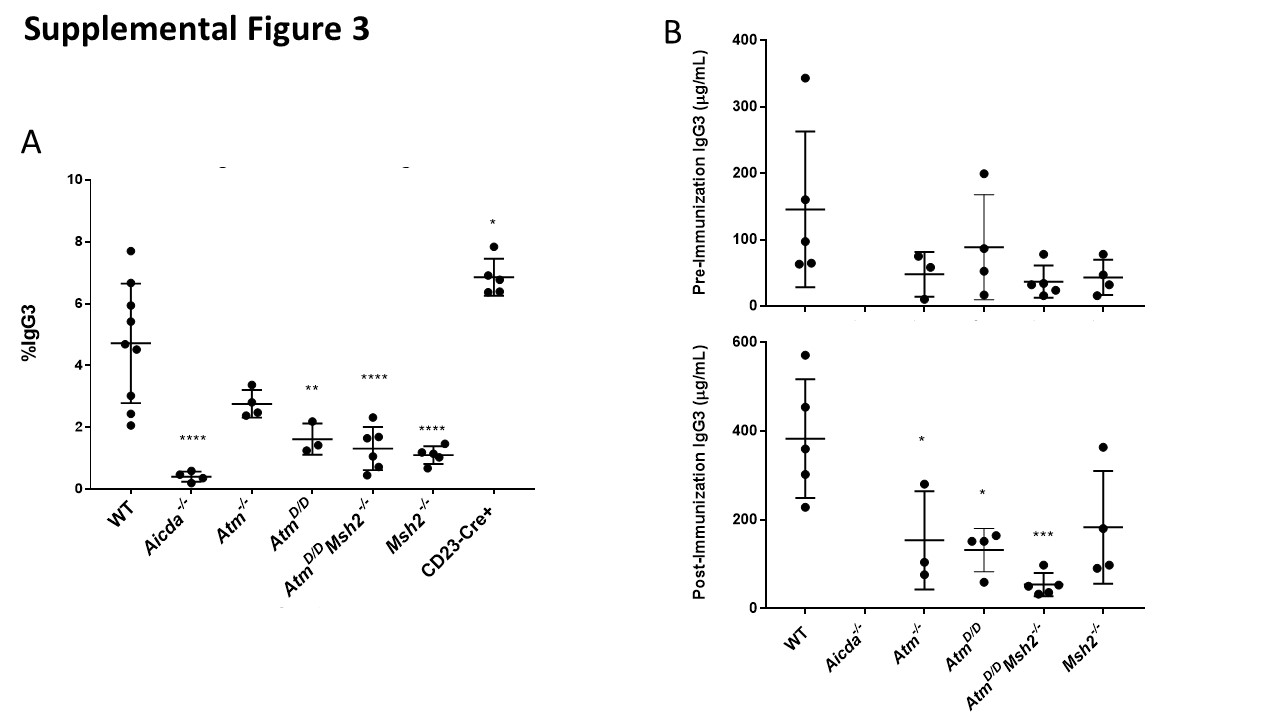

### Supplemental figure 5

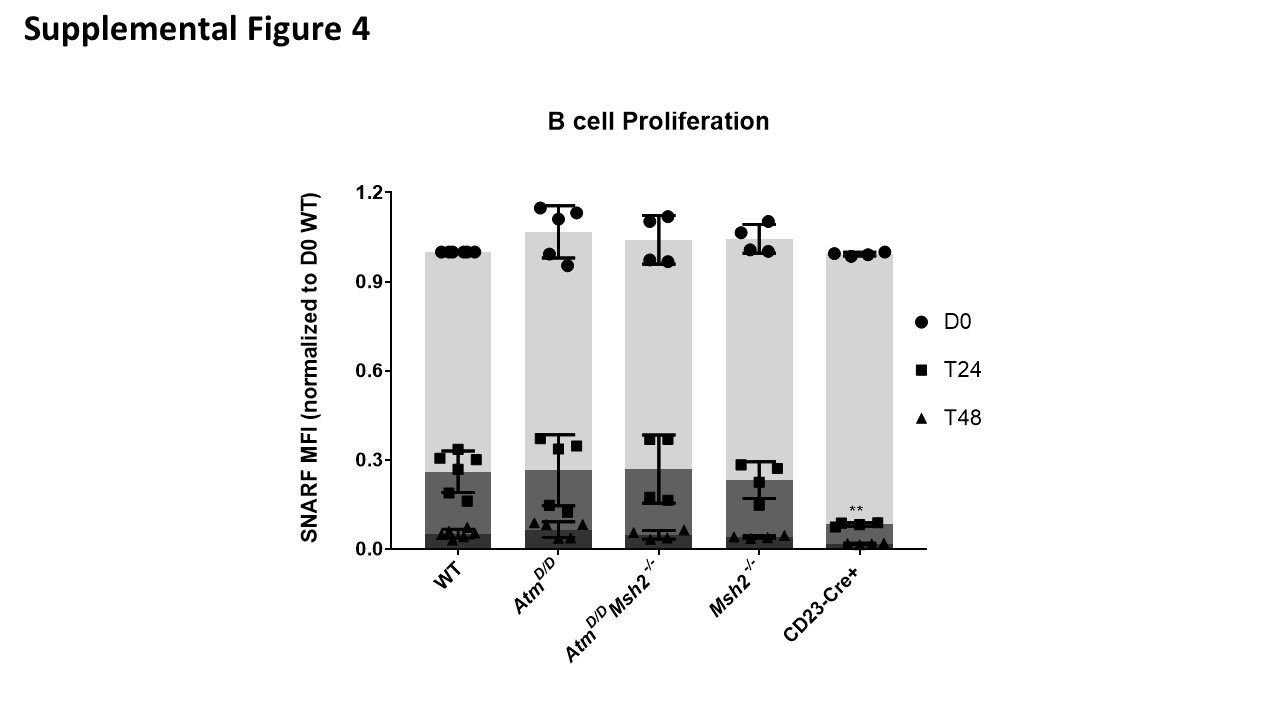

### Supplemental figure 6

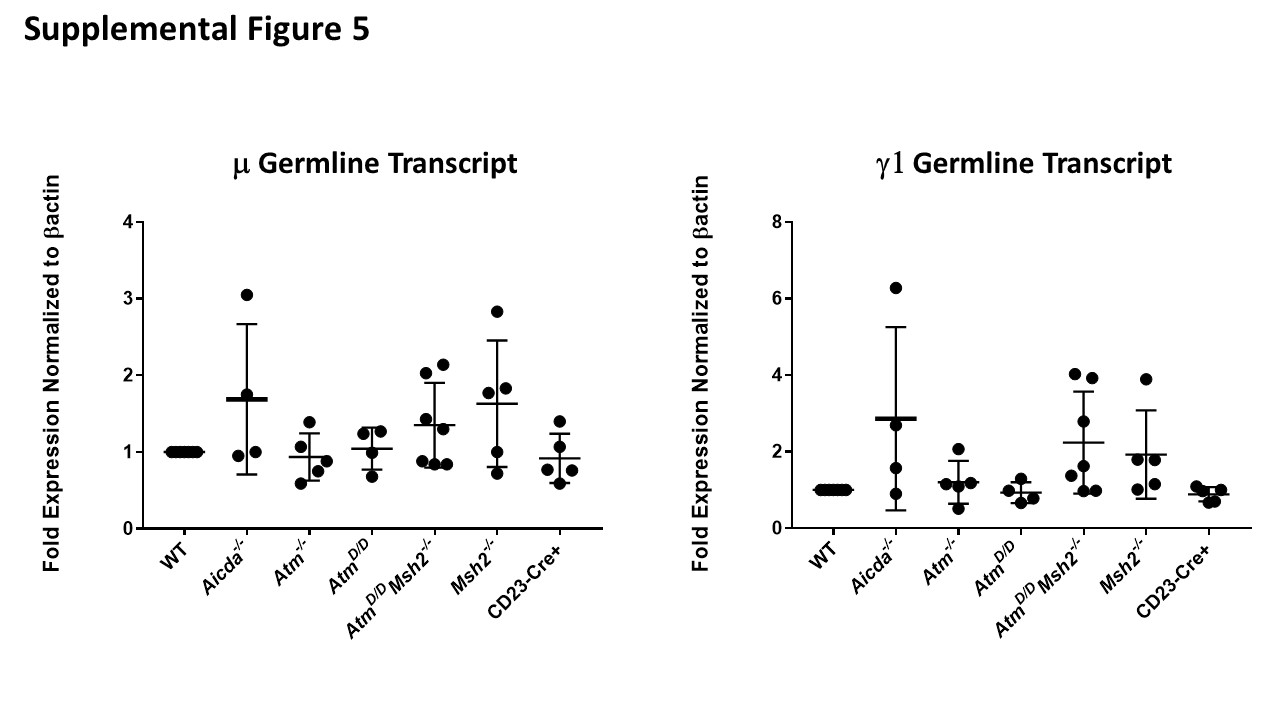
